# Divergent Evolutionary Pathways of Myxoma Virus in Australia: Virulence Phenotypes in Susceptible and Partially Resistant Rabbits Indicate Possible Selection for Transmissibility

**DOI:** 10.1101/2022.06.02.494607

**Authors:** Peter J Kerr, Isabella M Cattadori, Derek Sim, June Liu, Edward C. Holmes, Andrew F. Read

## Abstract

To characterize the ongoing evolution of myxoma virus in Australian rabbits we used experimental infections of laboratory rabbits to determine the virulence and disease phenotypes of recent virus isolates. The viruses, collected between 2012-2015, fell into three lineages, one of which, lineage c, experienced a punctuated increase in evolutionary rate. All viruses were capable of causing acute death with aspects of neutropenic septicaemia, characterized by minimal signs of myxomatosis, the occurrence of pulmonary oedema and bacteria invasions throughout internal organs, but with no inflammatory response. For the viruses of highest virulence all rabbits usually died at this point. In more attenuated viruses some rabbits died acutely while others developed an amyxomatous phenotype. Rabbits that survived for longer periods developed greatly swollen cutaneous tissues with very high virus titres. This was particularly true of lineage c viruses. Unexpectedly, we identified a line of laboratory rabbits with some innate resistance to myxomatosis and used these in direct comparisons with the fully susceptible rabbit line. Importantly, the same disease phenotype occurred in both susceptible and resistant rabbits, although virulence was shifted towards more attenuated grades in resistant animals. We propose that selection against inflammation at cutaneous sites prolongs virus replication and enhances transmission, leading to the amyxomatous phenotype. In some virus backgrounds this creates an immunosuppressive state that predisposes to high virulence and acute death. The alterations in disease pathogenesis, particularly the overwhelming bacterial invasions that characterize the modern viruses, suggest that their virulence grades are not directly comparable with earlier studies.

**IMPORTANCE:** The evolution of the myxoma virus (MYXV) following its release as a biological control for European rabbits in Australia is the textbook example of the co-evolution of virus virulence and host resistance. However, most of our knowledge of MYXV evolution only covers the first few decades of its spread in Australia and often with little direct connection between how changes in virus phenotype relate to those in the underlying virus genotype. By conducting detailed experimental infections of recent isolates of MYXV in different lines of laboratory rabbits we examined the ongoing evolution of MYXV disease phenotypes. Our results reveal a wide range of phenotypes, including an amyxomatous type as well as the impact of invasive bacteria, that in part depended on the level of rabbit host resistance. These results provide a unique insight into the complex virus and host factors that combine to shape disease phenotype and viral evolution.

---

The evolution of the myxoma virus after it was released in 1950 to control European rabbits (*Oryctolagus cuniculus*) in Australia is the canonical example of viral and host adaptation after a pathogen spillover (1, 2) (3, 4). By experimentally infecting laboratory rabbits with viral isolates collected over time in the field, evolved changes in the virulence and pathology caused by the virus can be determined (5). Most famously these studies demonstrated the initially highly virulent progenitor (99.8% case fatality rate CFR (6)) rapidly evolved into somewhat attenuated viral variants (CFRs 60-95%) that came to dominate in the field. Rabbits infected with these variants survived for longer, prolonging the infectious period giving these variants a transmission advantage (7). This remains the textbook example of the evolution of reduced virulence. However, the virus continued to evolve, and we recently reported that Australian viral isolates collected in the 1990s did not presented as classical myxomatosis in laboratory rabbits (8). Instead, they induced a novel amyxomatous disease phenotype that was characterized by a highly lethal immune collapse syndrome, consistent with massive immunosuppression. We interpreted this phenotype as an adaptation to overcome evolving resistance in the wild rabbit population (9, 10), a classic example of a biological arms race. Notably, some of these viruses killed laboratory rabbits more quickly than the progenitor strain, but this increase in virulence over the original progenitor is via very different pathways. A similar novel disease was seen with recent virus isolates from Great Britain suggesting convergent evolution (11).

Myxoma virus is a double-stranded DNA virus in the genus *Leporipoxvirus* (*Poxviridae*). The virus likely originated in the Americas in rabbit-like *Sylvilagus* species of lagomorphs. Although apparently largely innocuous in its natural hosts (12) (13), in the European rabbit MYXV causes the lethal, generalized disease myxomatosis. An isolate of MYXV, originally from Brazil in around 1910 (the Standard laboratory strain; SLS) (14) was experimentally released in Australia in 1950 to determine its potential as a biological control for the introduced European rabbit. The experiment exceeded all expectations with the virus spreading across large parts of south-eastern Australia and killing tens if not hundreds of millions of rabbits within a few months (15). In 1952, a landholder in France inoculated two wild rabbits with a separate Brazilian isolate of MYXV from 1949 (Lausanne strain; Lu). The virus spread from that introduction into wild and farmed rabbits across Europe (1) (16). In both continents, MYXV is now a permanent part of the rabbit’s environment.

The original Brazilian mammalian host species was not present in either Australia or Europe so there was no reservoir host to maintain a pool of unadapted virulent virus. The subsequent evolutionary outcomes for virus and rabbits were remarkably similar on both continents with natural selection for moderately attenuated viruses and for resistance in the rabbit population (3) (4). Because the laboratory rabbit is the same species as the wild European rabbit and the released progenitor viruses were available, the changes in both the virus and the wild rabbit could be followed using the progenitor virus and unselected laboratory rabbits as standards (17).

Most descriptions of clinical myxomatosis and disease pathogenesis are based on the progenitor viruses SLS and Lu or their close derivatives (1) (18) (3). Following intradermal inoculation MYXV replicates locally inducing hyperplasia, hypertrophy, and degeneration of epidermal cells together with massive disruption of the dermis and small blood vessels and deposition of large amounts of mucinous material. This primary lesion can develop to 3 to 5 cm in diameter and a few cm in height. Within 48 hours, virus can be detected in the draining lymph node, where it replicates to high titres and induces lymphocyte destruction and proliferation of stromal cells (19). Generalized infection follows, probably spread by infected lymphocytes (20).

The obvious clinical manifestations of myxomatosis caused by SLS and Lu are due to virus replication in cutaneous and mucocutaneous sites such as eyelids, nasal passages, base of the ears and anogenital region. This causes swollen heads, hard, swollen eyelids with copious mucoid to mucopurulent discharge together with swollen, drooping ears and upper respiratory tract occlusion, often with mucopurulent discharge from the nostrils, and massively swollen, red/black anogenital region; in males, scrotal oedema is prominent. Virus localization in the skin leads to discrete secondary lesions over legs, head, ears, and body, starting from around 6 days after infection. The virus is profoundly immunosuppressive, encoding multiple proteins that suppress or subvert the host immune response (3) (18). Death typically occurs 10 to 15 days after infection. The proximate cause of death is obscure with only limited pathology in key organs (21). Secondary bacterial infection of eyelids and nasal passages with gram negative bacteria such as *Pasteurella multocida* and *Bordetella bronchiseptica* is common but is generally not present in internal tissues of rabbits infected with highly virulent Lu or SLS (22) (23) (24). More attenuated viruses have a prolonged disease course with some rabbits able to control the virus and recover.

Transmission of MYXV is predominantly by mosquitoes or fleas that probe through the virus-rich epidermis of the cutaneous lesions, pick-up virus particles on their mouthparts and passively inoculate these at a subsequent feeding attempt on another rabbit. MYXV does not replicate within the vector and virus ingested as part of a blood meal is not transmitted (25). Efficient vector transmission depends on high titres of virus in cutaneous sites and rabbits surviving for sufficient time to transmit virus (7).

SLS killed rabbits in around 10 to 12 days, providing a relatively short window for transmission (6) (7). It quickly became apparent that slightly attenuated viruses were replacing the virulent virus in the field (26). Although they still caused very high CFRs, these viruses allowed infected rabbits to survive for longer in an infectious state and so had a higher probability of transmission and thus could be considered as fitter (7). To systematize these observations, virulence was measured by inoculating a low dose of virus into small groups of laboratory rabbits and calculating average survival times (AST) and case fatality rates. Virus isolates were classified into 5 virulence grades (Table 1) (5). Grade I virulence represented the originally released virus with CFR of essentially 100% and AST of ≤13 days, while grade V viruses killed fewer than 50% of infected rabbits. In later work, grade III viruses were subdivided into grade IIIA and grade IIIB (27). Between 1951 and 1980, most viruses were classed as grade III with CFRs of 60-95% and AST of 17-28 days. Hundreds of isolates were classified using this system in Australia and Britain and smaller numbers in other European countries.

**TABLE 1.** Virulence grades.

| Grade | AST<sup>1</sup><br>(days) | CFR (%)<sup>2</sup> |
| --- | --- | --- |
| I | ≤ 13 | 100 |
| II | 14-16 | 95-99 |
| IIIA | 17-22 | 90-95 |
| IIIB | 23-28 | 70-90 |
| IV | 29-50 | 50-70 |
| V | n/a | < 50 |

<sup>1</sup>AST average survival time <sup>2</sup>CFR case fatality rate. n/a not applicable

Resistance to myxomatosis emerged quite rapidly in Australian rabbits although it was unevenly distributed across the landscape, being stronger in hot arid regions and weaker in wet cool areas (9) (10). Selection pressure would have been very high since most sick rabbits will be removed by predators. Resistance appeared more slowly in Britain but once emerged seems to have spread rapidly (28) (29). Resistance also emerged in Europe although systematic testing was more limited (16). Genetic studies have revealed that mutations selected in rabbit populations from Australia and Europe are remarkably consistent and many involve genes of the innate immune response although specific mechanisms have not been demonstrated (30). Experimentally, resistant rabbits still became infected but were able to control early virus replication in the draining lymph node indicating a role for enhanced innate antiviral responses that favoured subsequent adaptive immune system control of the virus (19). Resistance can be overcome by more virulent virus strains and by skewing the immune response towards Th2 (31) (32). This emerging resistance in wild rabbits during the 1950s may explain why grade III viruses were most prevalent in the field although experimentally grade IV viruses were most transmissible when measured in unselected laboratory rabbits (7).

Alterations in disease phenotype other than virulence also occurred. Australian field viruses evolved a flat primary lesion compared to the raised domed lesion caused by SLS (1). In Europe, viruses emerged that caused a disease described as amyxomatous because infected domestic rabbits did not develop the typical cutaneous nodular lesions. Laboratory rabbits infected with these viruses still developed swollen heads and eyelids and other external signs of myxomatosis (33). Recent testing of viruses isolated in the 1990s from Australia and 2009-2011 in the UK showed that many of these also exhibited an amyxomatous phenotype in laboratory rabbits with limited pathology at the primary site despite high titres of virus. In addition, these recent Australian and UK viruses had a distinct and novel disease phenotype typified by neutropenia, collapse and death, often with minimal signs of myxomatosis but with haemorrhage in multiple tissues, pulmonary oedema, and frequently, large masses of bacteria throughout the tissues without any apparent cellular inflammatory response visible histologically. This suggested the development of novel evolutionary pathways to enhanced virulence from moderately attenuated viruses in the face of evolving resistance in the rabbit population: a biological “arms race” (8) (11). Remarkably, given the convergence of phenotype and the selection of common mutations in the rabbit populations, there were almost no shared mutations between the Australian and UK viruses. This suggests that these large viral genomes, of around 160,000 bp, offer multiple pathways to the same phenotype.

Herein, we address two key questions. First, given the novel disease phenotype in viruses from the 1990s, has the disease phenotype caused by viruses isolated 13 to 16 years later continued to evolve? In particular, has the virus evolved to become even more immunosuppressive, as might be expected from a continuing evolutionary arms race? Second, genome sequencing has revealed three main genetic lineages of MYXV in Australia – denoted a, b and c - with one (lineage c) experiencing a rapid burst of evolution between 1996 and 2012 (34). We therefore asked whether there were phenotypic differences in disease caused by these viruses that might point to the action of adaptive evolution.

## RESULTS

### Trial 1: Susceptible Oak rabbits

The Australian sequences of MYXV obtained since 1990 can be placed into three genetic lineages, labelled a, b and c (Fig. 1B) (34). Note in the original paper these lineages were labelled A, B, C. To avoid confusion with disease category classification (A, B, C, D) we have used lower case letters here for the virus lineage (a, b, c). Five viruses isolated in 2012, three from lineage c and two from lineage b, were tested in the same manner and rabbit line (Oak) as previously described (8) (11). Survival plots and virulence grade assignments are shown in Fig. 2.

**FIG 1.**
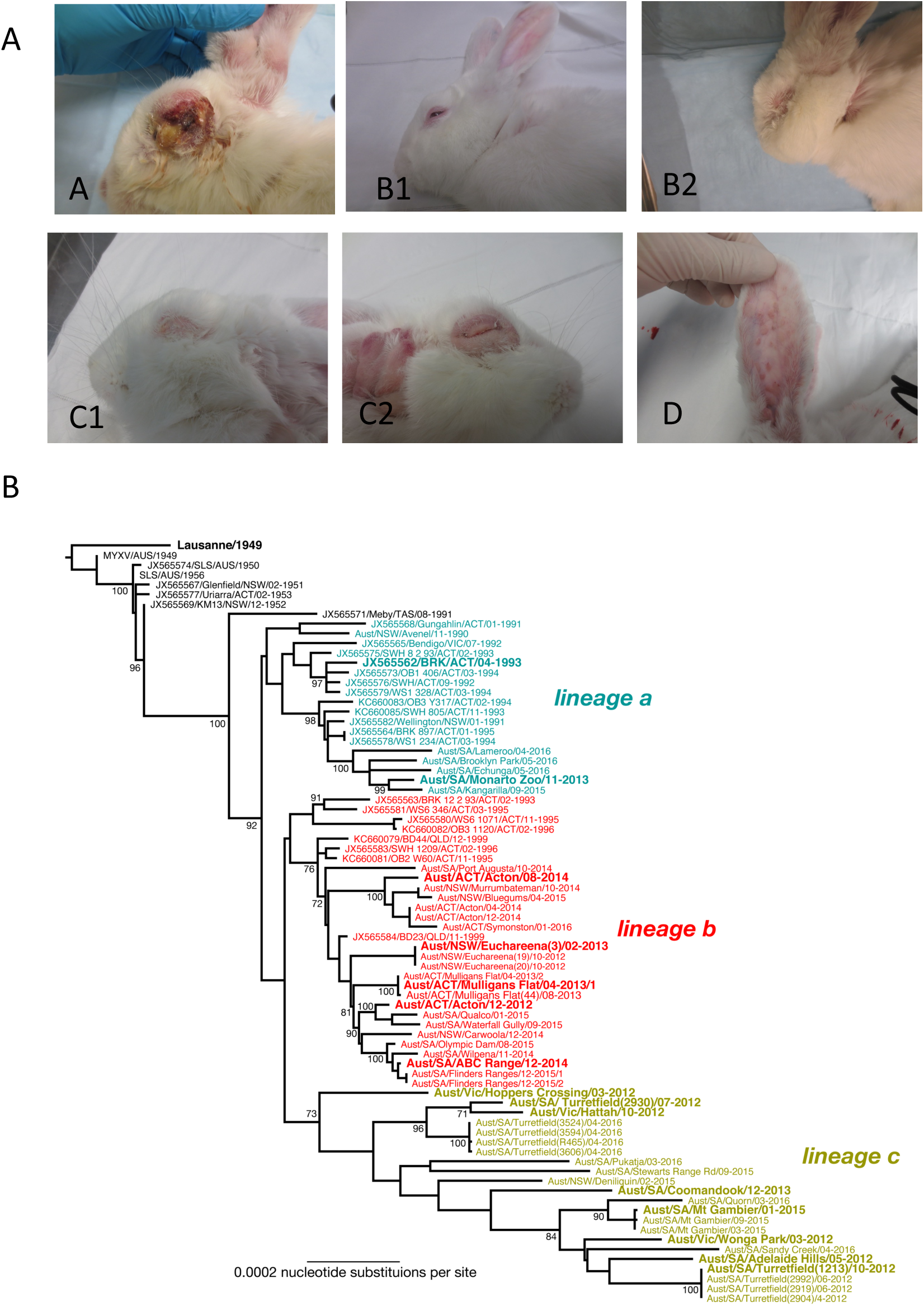
Disease phenotype categories. Phenotypes were allocated to categories A to D at time of death based on the criteria described in the methods. Category A: rabbit infected with Lu at day 12. For categories B and C two examples of each are shown to cover the range of phenotype. B1: ABC Range virus at day 8. B2: Adelaide Hills virus at day 11. C1: ABC Range virus day 19. C2: Wonga Park virus day 19. For category D the secondary lesions on the inside of the ear at day 20 are shown for a rabbit infected with Mulligans Flat virus that made a rapid recovery. (B) Phylogeny of MYXV in Australia. The tree shows the evolutionary relationships between 78 complete MYXV genomes with the three main lineages (a, b, c) highlighted in different colours. The viral isolates used in this study are shown in larger bold font. The tree is rooted on the Lu strain, with branches scaled according to the number of nucleotide substitutions per site. Bootstrap values are shown for representative nodes. The tree was redrawn from that presented in (34).

**FIG 2.**
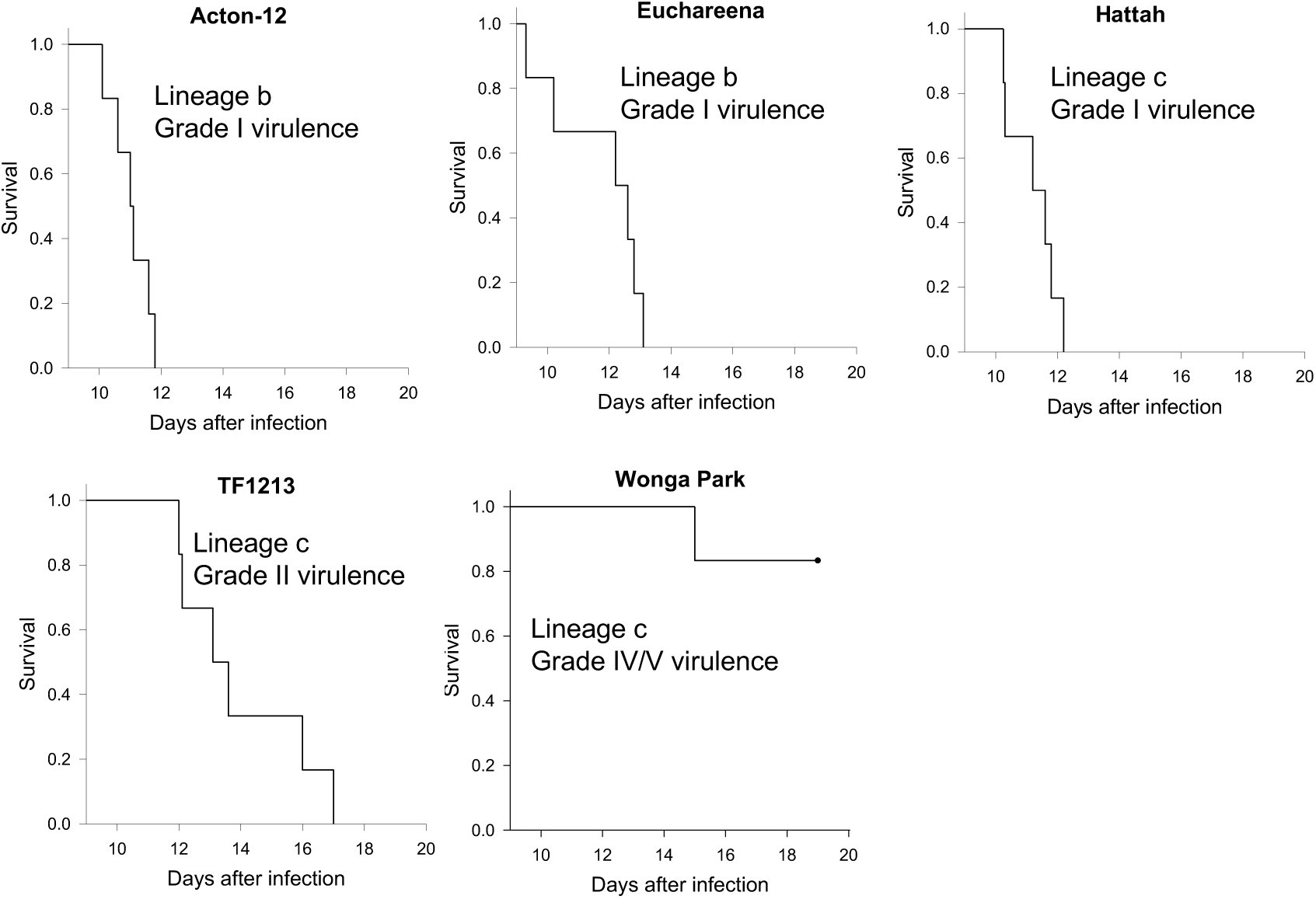
Survival plots and virulence grades for trial 1 (Oak rabbits).

Three of the viruses tested - Hattah (lineage c), Euchareena (lineage b), Acton-12 (lineage b) - had highly lethal grade I virulence, amyxomatous phenotypes (category B on the disease appearance scale) characterized by 100% case fatality with clinical signs and pathology typical of the highly immunosuppressive phenotype previously described (8) (11).

Clinically, rabbits infected with these three grade I viruses developed moderately swollen heads together with weight gain between days 7 to 10 suggesting tissue oedema. Eyelids tended to be moderately swollen with eyes still open at time of death. Ears were generally only swollen at the base and there was usually some degree of ano-genital swelling. Primary lesions were flat, diffuse and poorly defined from the surrounding skin with no scabbing. Rectal temperatures transiently spiked around day 4 to 5 then returned to normal but most rabbits had a second high spike to as high as 42°C in the 24 hours prior to death. All rabbits died or were euthanised between days 9 and 13 after infection. In some cases, rabbits rapidly became moribund having been apparently normal several hours earlier. Others were found with respiratory distress or trembling and recumbent. At necropsy, there was typically massive pulmonary oedema; histopathology often revealed bacterial invasion of multiple tissues such as lung, lymph nodes, spleen, heart, liver and kidney. The primary lesions ranged in thickness on cut section from 1 to 5 mm with most 1 to 2 mm. Often there was subcutaneous oedema fluid over the rump and down the hind legs.

Interestingly, although the Acton-12 isolate showed an amyxomatous acute death phenotype in laboratory rabbits, the wild rabbit from which it was isolated clearly had multiple flat lesions of the typical myxomatous phenotype (Supplementary material, Fig. S1a,b). TF1213 (lineage c) was still highly lethal (grade II virulence), with four of the rabbits exhibiting pulmonary oedema/toxaemia around days 12 to 13, and three of these having a temperature spike in the 24 hours prior to death. The remaining two rabbits were euthanized on day 15 but one had pulmonary oedema on histology and the other had bacteria in tissues indicating a similar pathogenesis to the more acute deaths. Compared to the lineage b viruses rabbits infected with TF1213 had an obviously more proliferative disease with more swelling around head, eyelids and ears and oedematous ridging of the skin of the anterior surface of the forelegs. There was some hair loss over the head area. Eyelids were fully closed at time of death. Ano-genital swelling was much more severe and there was usually some scrotal oedema. Primary lesions were more defined, slightly raised and 3 to 10 mm thick on cut sections. Subcutaneous oedema of rump and hind legs was also a feature.

Wonga Park (lineage c) was quite different with only one rabbit moribund and euthanized at 15 days with haemorrhage and pulmonary oedema. Three of five of the remaining rabbits developed extreme swelling of head and ears with marked increase in body weight while the other two had a less severe clinical progression and also developed multiple coalescing secondary lesions. A feature of the Oak rabbits infected with this virus was grossly swollen, drooping ears and extreme folding of the thickened skin at the base of the ears and over the face (Supplementary material, Fig. S1c,d). Swollen drooping ears are often seen with MYXV isolates from the 1950s but were rarely seen with the more recently isolated viruses we have tested. All five remaining rabbits were euthanized on day 19 and scored as category C on the disease appearance scale but these had much more cutaneous swelling than we had previously seen. It was difficult to predict the outcomes for these rabbits and so define an endpoint for euthanasia, but the virus was obviously considerably attenuated and so continuing the trial could not be justified; at day 19, when the trial was terminated, the surviving rabbits were all still eating and drinking.

### Trial 2: Partially resistant CR rabbits

At this point in our investigation, a change in rabbit supplier occurred. This meant that NZ White rabbits from an independent genetic line were supplied – herein named CR rabbits. To control for any variation between the rabbit lines, we tested a previously well characterized, highly virulent lineage a virus, BRK isolated in 1993, along with nine viruses isolated between 2012 and 2015 from lineages a, b and c not previously tested. Based on experience with multiple rabbit lines, we were not anticipating any differences between the lines of NZ White rabbits. However, it quickly became obvious that this CR line of rabbits had some innate resistance to myxomatosis compared to previously tested laboratory rabbits in the USA and in Australia.

The control virus, BRK, had previously been shown to have grade I virulence in Australia and in multiple trials in the US (35) (8). Infection was characterized by sudden death between 11-13 days after inoculation typically with pulmonary oedema/toxaemia and an amyxomatous phenotype. However, in the CR rabbits, no deaths occurred before day 13 and one rabbit had a prolonged survival and was euthanized at day 24. Acute deaths with pulmonary oedema/toxaemia still occurred in four of the rabbits (Fig 3). Interestingly, this circa IIIA virulence grade (AST 16.5 days puts it on the line between grade II and IIIA) was very similar to results obtained in a previous experiment when Oak rabbits infected with BRK were prophylactically treated with a broad spectrum antibacterial during the clinical phase of the disease. This was intended to reduce the effects of bacterial infection likely associated with neutropenia and AST was 16.1 days compared to 11.7 days for rabbits without antibiotic treatment (8).

**FIG 3.**
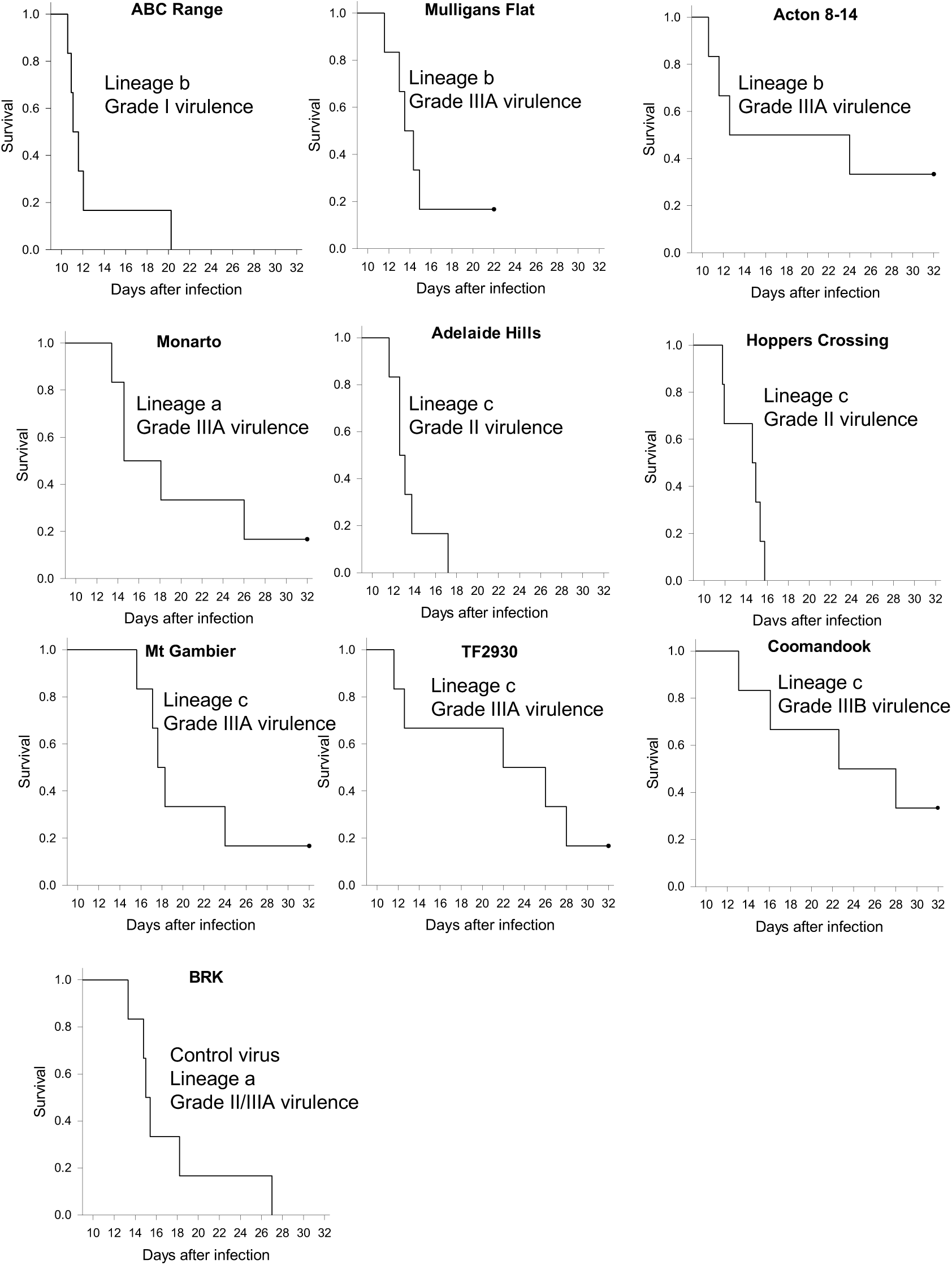
Survival plots and virulence grades for trial 2 (CR rabbits).

The nine new viruses tested were mostly of grade II or III virulence in the CR rabbits with only ABC Range being grade I. Prolonged survival of one or more rabbits was a feature of all the viruses tested in the CR rabbits (Fig. 3). However, it was clear that the previously described novel disease phenotype still occurred in these CR rabbits with deaths characterized by pulmonary oedema and bacteria through multiple tissues occurring in all groups (see below). Many longer-term survivors developed pneumonia, and some had severe pathology of testes and epididymis including caseous necrosis. Most of these were euthanized on humanitarian grounds rather than being allowed to die (across the 3 trials 73% of rabbits were euthanised). Overall, there was a wider range of survival times between rabbits for each virus tested in the CR rabbits compared to earlier results in the Oak rabbits. Even the most virulent virus tested, ABC Range, had one rabbit with more prolonged survival (Fig 3).

With the exception of seven rabbits (see below) all of the viruses generally caused an amyxomatous phenotype with minimal primary lesions and a lack of secondary lesions. All viruses caused some acute deaths due to pulmonary oedema/toxaemia, where these occurred on or before day 15 they were classed as category B. Those rabbits that survived longer, including many scored as recovered or likely to recover, were scored as category C. A few of these rabbits developed secondary lesions mostly around head and ears. Mt Gambier caused death with pulmonary oedema and bacteria in tissues, but these deaths occurred after day 15 and so were somewhat arbitrarily classed as category C. It was difficult to categorize four rabbits infected with TF2930, that developed secondary lesions on the ears between days 10 and 12, two of which developed more generalized secondary lesions. In addition, three rabbits (one from the Mulligans Flat group and two from the Acton-14 group) developed secondary lesions early and had a mild disease course with rapid recovery. These seven rabbits have been classed as category D (attenuated myxomatous disease) based on the secondary lesion development, but the disease course could be considered atypical since the primary lesion was clearly of the amyxomatous type.

Rabbits in category C infected with lineage c viruses that survived for longer tended to develop the extreme folding of the thickened skin of the base of the ears and anterior skin of the forelegs seen in Wonga Park in the previous trial, this was particularly the case with the attenuated Mt Gambier and Coomandook viruses. However, one rabbit infected with the lineage a Monarto virus that was euthanized at day 26 developed very similar lesions. The assignment of a virulence grade for Acton 8-14 is somewhat arbitrary since with 2 survivors, both of which had an atypical disease course, it could be considered a grade IIIB or IV virus but on AST of 19.1 days was a grade IIIA virus. Similarly, TF2930 on survival times was grade IIIA virulence virus but this was skewed by the two acute deaths with the remaining rabbits having much more prolonged survival more like a grade IIIB or IV virus.

### Trial 3: Direct comparison of susceptible and partially resistant rabbits

To further investigate this observed resistance in the CR rabbits, we obtained Oak rabbits from the previous supplier (although ownership had changed) and CR rabbits to undertake a simultaneous comparison of infections between the two lines of rabbits. For this we selected eight viruses across a range of virulence grades: six used in trial 2 in CR rabbits, Wonga Park from trial 1 as a highly attenuated virus, and the Lausanne strain as a known grade 1 virulent virus with the classic myxomatous nodular phenotype.

Note survival time results and some disease data for Lu in the susceptible (Oak) rabbits from this trial have been included in an earlier paper (11) and are here included only for comparison with the CR rabbits and with the cutaneous lesion morphology and inflammatory response and titres of the Australian viruses.

The virulence grades resulting from this trial are summarized in Table 3 and the AST and assigned virulence grades for trial 2 are also shown here for comparison; Fig 4 shows the survival analyses for the trial. In general, the viruses are more attenuated in the CR rabbits. The most obvious differences were seen between the two groups of rabbits infected with the highly virulent Lu and ABC Range viruses and with the very attenuated Wonga Park virus. CR rabbits infected with Lu always showed less severe clinical signs compared to the Oak rabbits infected with Lu at the same timepoints. This was easily seen because of the very florid nature of the clinical signs associated with Lu infection and was obvious to independent observers who were unaware of the different groups of rabbits. Deaths in the CR rabbits did not commence until all the Oak rabbits were dead. Using the Fenner method for calculating AST, virulence was graded as I for Oak rabbits and II for CR and the difference in the Kapler-Meier survival analysis was significant (p = 0.018) (Fig. 4). Subsequent experience with the Lu virus in CR rabbits has confirmed this prolonged survival time and grade II virulence (Cattadori unpublished data).

**FIG 4.**
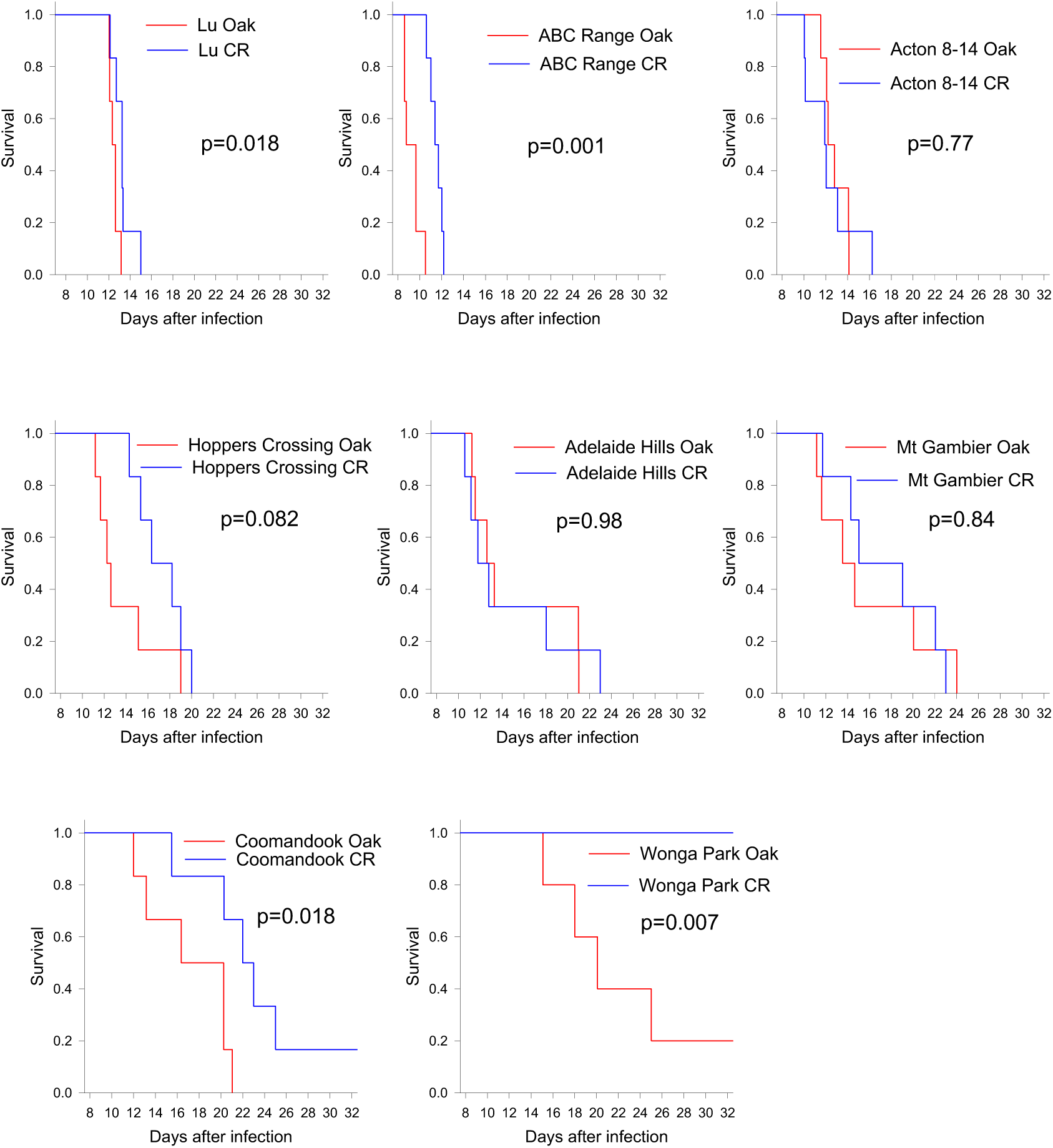
Comparative survival plots for Oak and CR rabbits (Trial 3). For each virus tested the survival of Oak and CR rabbits is shown on the same plot. P values show significance of difference between the plots (log rank tests).

No clinical differences were seen between the two groups of rabbits infected with ABC Range; both groups had an amyxomatous acute death phenotype category B (see Fig. 5B below). ABC Range was of grade I virulence in both groups of rabbits but, similarly to Lu, all of the Oak rabbits were dead when deaths commenced in the CR rabbits and the difference in the survival time was significant (p = 0.001; Fig. 4).

**FIG 5.**
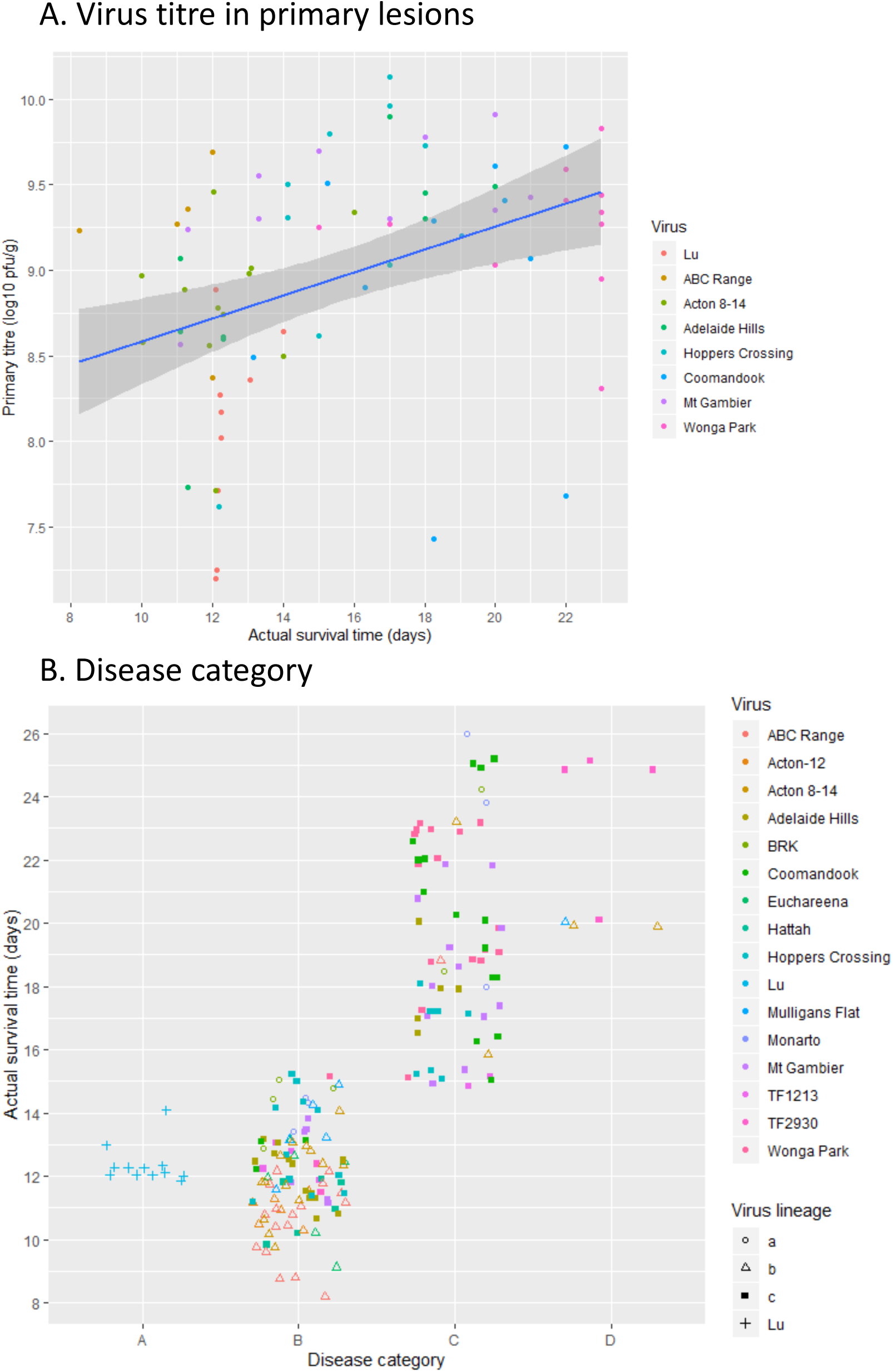
Primary lesion titres and disease phenotype categories. (A) Titres in primary lesions from trial 3. Scatterplot of virus titre at actual time of death and regression line with 95% confidence intervals. (B) Disease phenotype scores. Each rabbit was assigned a disease category (A, B, C, D) at actual time of death. Points are coded by virus lineage, a, b or c for the Australian viruses and Lu. Points have been jittered 0.3 units horizontally and 0.25 vertically for clarity.

At the other end of the virulence spectrum, the highly attenuated Wonga Park also showed clear differences in virulence, survival curves and clinical signs between the two groups of rabbits. In Oak rabbits, Wonga Park was estimated as grade IIIB virulence, whereas it was assigned grade V virulence in the CR rabbits (Table 3). Clinical signs were generally less severe in the CR rabbits although both groups exhibited an amyxomatous category C clinical appearance. One rabbit in each group developed some late secondary lesions. Secondary lesions mostly developed from the thickened, folded skin of the forelegs and base of the ears and were thus developmentally quite different to the early secondary lesions seen in accelerated recoveries in trial 2 or the widespread secondary lesions seen in rabbits infected with Lu. Drooping ears, seen in the Oak rabbits infected with Wonga Park did not occur in the CR rabbits. Survival times were significantly different between the 2 groups (p = 0.007; Fig. 4) with no rabbits euthanized or dead in the CR group prior to termination of the trial at day 23 whereas in the Oak group of rabbits, 4 rabbits were euthanized between days 15 and 22 and only one was scored as a likely survivor. Even with this very attenuated virus, the earliest death in the Oak rabbits (day 15) was associated with bacteria in multiple tissues as also seen in trial 1 in one rabbit.

Rabbits infected with Hoppers Crossing, Mt Gambier and Coomandook viruses showed similar trends between the groups with lower estimated virulence in the CR rabbits (Table 3) but no significant difference in the survival curves (Fig. 4). Acton 8-14 (grade I virulence in this trial) and Adelaide Hills (grade II virulence) viruses showed no differences between the 2 groups of rabbits.

Results from trial 2 with CR rabbits were not always duplicated in this comparative trial. This was particularly the case for Acton-8-14, which was a grade IIIA virulence in the previous trial but a grade I virulence in the comparative trial with no difference between the Oak and CR rabbits and an amyxomatous phenotype (Table 3). The wild rabbit from which this virus was isolated had a clearly myxomatous phenotype with grossly thickened eyelids and discrete secondary lesions on muzzle and at the base of the ear (data not shown). Inspection of the survival curve for trial 2 (Fig. 3) shows that the results were dominated by one rabbit which had a prolonged survival time and two with a rapid recovery, the time of death for the other three rabbits was similar to those in trial 3 (Fig. 4). A similar, although less pronounced, difference can be seen in the results for ABC Range, where in trial 2, one rabbit had a longer survival time. In trial 2, no rabbits infected with Mt Gambier were in disease category B. However, in this trial two of the CR rabbits and four of the Oak rabbits died with the acute death, category B disease type. This is the sort of variation that might be expected when there is a degree of innate resistance to MYXV in the rabbit population. It also indicates that the Fenner virulence scoring system using AST in small groups of rabbits breaks down with the modern viruses, where stochastic factors influencing severity of neutropenia and opportunistic bacterial invasion may influence the acute death phenotype such that moderately attenuated viruses could easily be scored as grade I on the basis of a single trial.

Some degree of innate resistance as seen here with the CR rabbits will exacerbate this problem with reproducibility by increasing the variation between rabbits.

### Virus titres

Virus titres were measured in spleen, popliteal lymph node, lung, liver and primary lesion collected at autopsy in trial 3. Because of the range of survival times when samples were taken and considering that rabbits that died overnight were not sampled, these findings are not as informative as would be the case if all animals were sampled at the same time points, making detection of differences in titres between rabbit lines unlikely. As there was no significant difference in primary lesion titres demonstrated between Oak and CR rabbits for any of the viruses (rank sum test) the titres for each virus at each time point have been pooled. Despite these limitations, some interesting trends can be seen. Titres in cutaneous lesions are of interest because these correlate with transmissibility (7). Primary lesion titres were very high at all times (Fig. 5A; Supplemental material Fig. S3e) and were clearly lower in the rabbits infected with Lu but otherwise there were no clear trends across time or viruses. The absence of any significant difference in primary lesion titres between the Oak and CR rabbits is compatible with earlier measurements in resistant wild rabbits where titres in the primary lesion were similar in resistant and susceptible rabbits (19). Virus titres were also measured in eyelids, the very thickened folded skin of the ears and discrete secondary lesions in rabbits infected with Coomandook and Wonga Park at late stages and were generally >10^9^ pfu/g in both groups of rabbits. These are extremely high titres; 10^7^ rabbit ID_50_/g is regarded as the threshold for high mosquito transmissibility (7). Virus titres have also been pooled for CR and Oak rabbits for the other tissues examined. Titres in lymph node tended to be fairly steady over time although some rabbits had much lower titres than others. Similarly, spleen titres tended to remain elevated but with a very large scatter. Lung and particularly liver titres declined with time (Supplemental material Fig. S2, Fig. S3).

Previous time course studies suggest that titres peak in lung and liver between 8 to 10 days after infection (8) and so are poorly represented in this data set. The rabbits infected with Lu stand out as tending to have lower titres in the tissues, particularly compared to virulent Australian viruses, even though this is a highly virulent virus (Supplemental material Fig. S3). These boxplots do not take time into account and so Lu is being compared with viruses such as Coomandook, Mt Gambier and Wonga Park that are attenuated with longer survival times and lower titres in liver and lungs at the much later times of death. This is similar to previous time course studies where the lineage a virus BRK had much higher titres in liver and lung than the progenitor SLS virus (8). The trend to lower titres of virus in the Lu rabbits supports the alteration in disease pathogenesis seen with the recent virus isolates.

### Clinical disease category

Fig. 5B summarizes the clinical disease category for all rabbits in the three trials. Only Lu consistently caused typical nodular myxomatosis (category A). Most rabbits infected with Australian viruses had the acute death phenotype, category B (100 of 173). However, lineage c viruses tended to have a more swollen head and eyelids than the lineage b viruses within this category (Fig. 1A category B2). Within category C there was a clear difference in appearance between those rabbits that survived for longer from lineage b and those from lineage c with the lineage c viruses causing much more proliferative, and folded cutaneous swelling and much more swelling and hardening of the eyelids (Fi.g 1A category C2). However, the lineage c viruses were generally more attenuated than the lineage b viruses and so had more rabbits surviving for longer plus we tested more lineage c viruses across the three trials and in the resistant CR rabbits. Longer survival times were also associated with CR rabbits compared to Oak. Only a single new lineage a virus was tested (Monarto) but results have also been included for BRK used as a control virus in trial 2. One rabbit infected with Monarto that was euthanized at 26 days (considered a likely survivor) had the deeply folded proliferative ear and head swelling that occurred in the attenuated lineage c viruses as did one rabbit infected with BRK euthanized at day 24 and considered very unlikely to survive. TF2390 from lineage c was somewhat ambiguous as two rabbits died acutely with pulmonary oedema (disease category B), but the others developed secondary lesions on the ears between days 10 and 12 and had prolonged survival times. However, the primary lesion was typical of the amyxomatous viruses. Somewhat arbitrarily these were classified into category D. Only three other rabbits were classed as category D (attenuated nodular myxomatosis). All three were CR rabbits infected with lineage b viruses that developed widespread secondary lesions early and had accelerated recoveries and primary lesions typical of amyxomatous disease rather than the attenuated myxomatous phenotype.

### Pathology

In general, necropsy findings in all three trials recapitulated our findings reported for Australian viruses from the 1990s and some recent UK viruses (8) (11). Two different syndromes were seen with the Australian viruses: (i) acute death early in the disease course typically with signs of toxaemia such as pulmonary oedema; (ii) more prolonged survival, gradual deterioration, sometimes with bacterial or viral pneumonia but often no obvious cause of death. In many cases euthanasia was necessary on welfare grounds. Rabbits infected with Lu had a very different florid nodular myxomatous disease with rapid onset of very swollen, hard eyelids, large raised, demarcated and scabbed primary lesions, very swollen ano-genital regions and swollen lips and ears plus raised discrete secondary lesions over the body (disease category A). The distinct difference between Lu and Australian isolate primary lesions is shown in Supp Data Fig 1E, Wonga Park day 22 primary lesion is pink, flat, poorly differentiated from surrounding tissue and shows no evidence of scabbing whereas at day 10 in the Lu primary lesion (Supplemental material Fig. 1F) the raised, demarcated, purple primary lesion is already starting to scab.

For the Australian viruses, rapid collapse and death between days 8-15 occurred unpredictably with relatively minor clinical signs of myxomatosis and was most often due to acute pulmonary oedema with massively swollen lungs and fluid and froth filling the bronchi and trachea (Fig. 6A). Sometimes haemorrhagic muscles and popliteal lymph node in one leg were evidence of bacterial invasion. Histologically, bacteria were often present in multiple tissues including heart, lung, lymph nodes, spleen, kidney, testis and liver. The presence of bacteria in the hepatic portal veins and sinusoids (Fig. 6C) suggests possible spread from gut tissues. A focal or generalized necrotic appendicitis with necrosis of lymphocytes and bacterial invasion also occurred (Fig. 6B and 6D). There was no evidence of cellular inflammatory responses around the bacteria in cases of acute death. Lymphoid tissues (regional lymph nodes, appendix, spleen, thymus) were generally devoid of lymphocytes or, in the thymus and appendix, contained only necrotic lymphocytes. Early degeneration of the seminiferous epithelia of the tubules of the testes was generally present. Occasional animals died during this time period with only minimal pathology and no bacteria visible in the internal tissues examined histologically.

**FIG 6.**
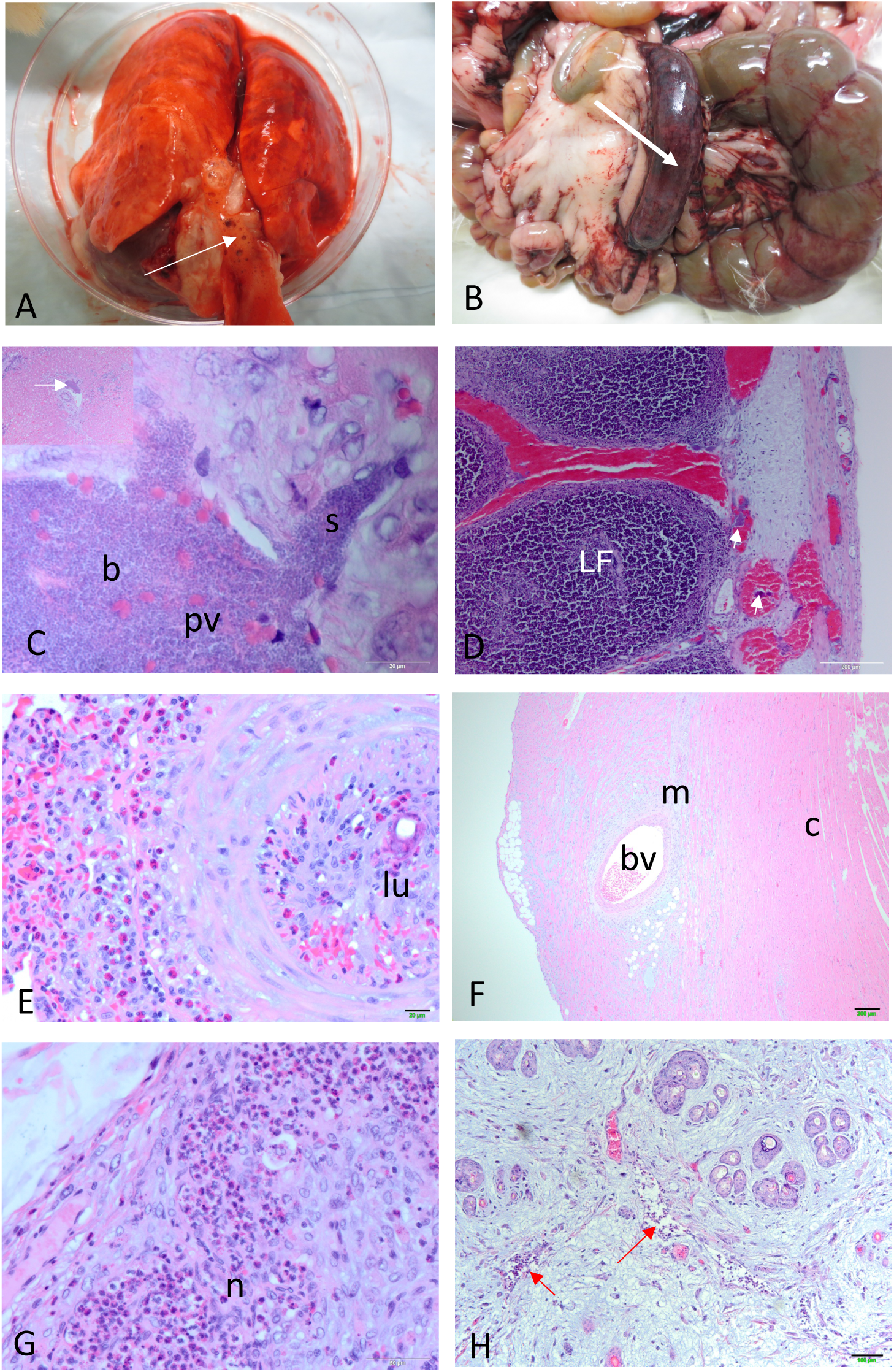
Pathology. (A) Pulmonary oedema. Swollen, fluid filled lungs with fluid and froth filling the trachea (arrowed). (B) Generalized swelling and haemorrhage in appendix (arrowed). (C) Bacteria (b) within a portal vein (pv) of the liver and spilling out into a sinusoid (s) (oil immersion). Inset top left shows lower magnification image with arrow indicating the bacteria within the same portal vein. (D) Histological section of inflamed appendix (lumen is on the left hand side of the figure). Lymphoid follicles (LF) are full of necrotic lymphocytes and surrounded by red blood cells within sinuses. Bacteria are present within sinuses and blood vessels (small arrows). (E) Obliteration of lumen of pulmonary blood vessel with neutrophil (heterophile infiltration); lu lumen of pulmonary blood vessel; Coomandook day 26. (F) Mucin-like material within the ventricle wall of the heart; bv blood vessel; c cardiac muscle; m mucin-like material (Hoppers Crossing day 15). (G) Lu popliteal lymph node. Neutrophil (n) (heterophile) influx. Note loss of lymphocytes (day 12). (H) Lu secondary lesion. Arrows indicate neutrophil influx within blood vessels. Note the mucin-like material dominating the section (day 12).

Early primary lesions were poorly differentiated from the surrounding skin. Histologically there was some hyperplasia and hypertrophy of epidermal cells, including the cells of the hair follicles, and minor disruption of the upper dermis evident as disruption of the collagen fibres together with deposition of grey-blue-staining mucin-like material. Occasional macrophages were present but there was a general absence of inflammatory cells such as neutrophils or lymphocytes. Fibroblasts were often prominent in the upper dermis (see below).

Despite the presence of bacteria in multiple tissues, neutrophils were absent from the tissues of rabbits that died acutely from infection with the Australian viruses. This was in contrast to rabbits that died at similar time points following infection with Lu, where bacteria were not present, but neutrophils were prominent in tissues such as primary and secondary lesions, lymph nodes, liver and spleen (Fig. 6G and 6H); (cytoplasmic granules in rabbit neutrophils stain with eosin and so appear red in histological sections and are often referred to as heterophiles).

Those rabbits that died or were euthanized at later time points generally had a different pathology. In some cases, it seems likely that the rabbits had survived an earlier bacterial invasion. In one rabbit infected with Coomandook clumps of bacteria were seen within the lung together with considerable inflammation. Death occurred at 14 days and was associated with pulmonary oedema, but the lung histology suggests that this probably is a rabbit that survived an initial bacteraemia long enough for bacterial invasion to occur and was able to mount some inflammatory response. Coomandook is an attenuated virus, and this was a CR rabbit. In contrast a CR rabbit infected with the same virus that died 24 hours earlier had generalized bacteraemia including within the heart muscle but without the inflammatory response. Other rabbits at later time points had interstitial pneumonia, which was probably due to the virus, and some also had secondary bacterial pneumonia but whether this stemmed from an earlier bacteraemia or a subsequent invasion was not clear. Histologically, at later time points, a proliferative obliteration of the lumen of large pulmonary blood vessels originally described by Hurst (1937) (24) was also observed together with neutrophil (heterophile) influx (Fig. 6E). Deposition of large amounts of mucinous material similar to that in the cutaneous lesions was often present in the connective tissue around blood vessels in multiple tissues including lung, heart, thymus and liver. Fig. 6F shows this in the cardiac muscle of the ventricle. Since this material is secreted in areas with active virus replication this suggests that there may be virus replication occurring in the heart and inducing some pathology. To our knowledge this has not previously been reported. Lymph nodes were generally depopulated often with proliferation of stromal cells but in some cases, a few lymphocytes could be seen migrating into the nodes. Damage that probably occurred during an earlier bacterial localization was seen in the kidney from a rabbit at day 24 with a mainly monocytic cell infiltrate and destruction of renal corpuscles in the cortex. Occasional rabbits showed mid-zonal degeneration of the liver (24). Pathology of the testis and epididymis could be very severe with both acute and chronic inflammation including caseous necrosis of the testis (data not shown).

The primary lesion histopathology of the Australian isolates ranged from relatively minor proliferation and hypertrophy of epidermal cells to ballooning degeneration of hypertrophied cells with vesicle formation and disruption of the upper and lower dermis and deposition of large amounts of blue-grey staining mucin-like material. Occasional eosinophilic cytoplasmic inclusions were seen in epidermal cells, but these were not common. The extent of the pathology varied with both survival time and virus lineage. Proliferation of the epidermal cells of the hair follicles was particularly prominent (Fig. 7). Inflammatory cell infiltration was minimal in primary lesions and only visible deep in the dermis if at all. Fibroblasts tended to be prominent through the disrupted dermis. Histologically, in the primary lesions the lineage c viruses tended to have more hyperplastic epidermis, which appeared to originate from the hair follicles and large amounts of mucinous material deposited in the dermis together with more disruption/swelling of the lower dermis compared to the lineage a and b viruses. Unlike in the Lu infected rabbits, scabbing of the primary and other cutaneous lesions was not a feature except very late in some eyelids, where a scab and accompanying inflammatory response occurred. In contrast, rabbits infected with Lu had deeply scabbed primary lesions (see also Supplemental material Fig. 1F) and eyelids at the time of death (day 12 to day 14) and some secondary lesions were also beginning to scab. Neutrophils were present as were macrophages.

**FIG 7.**
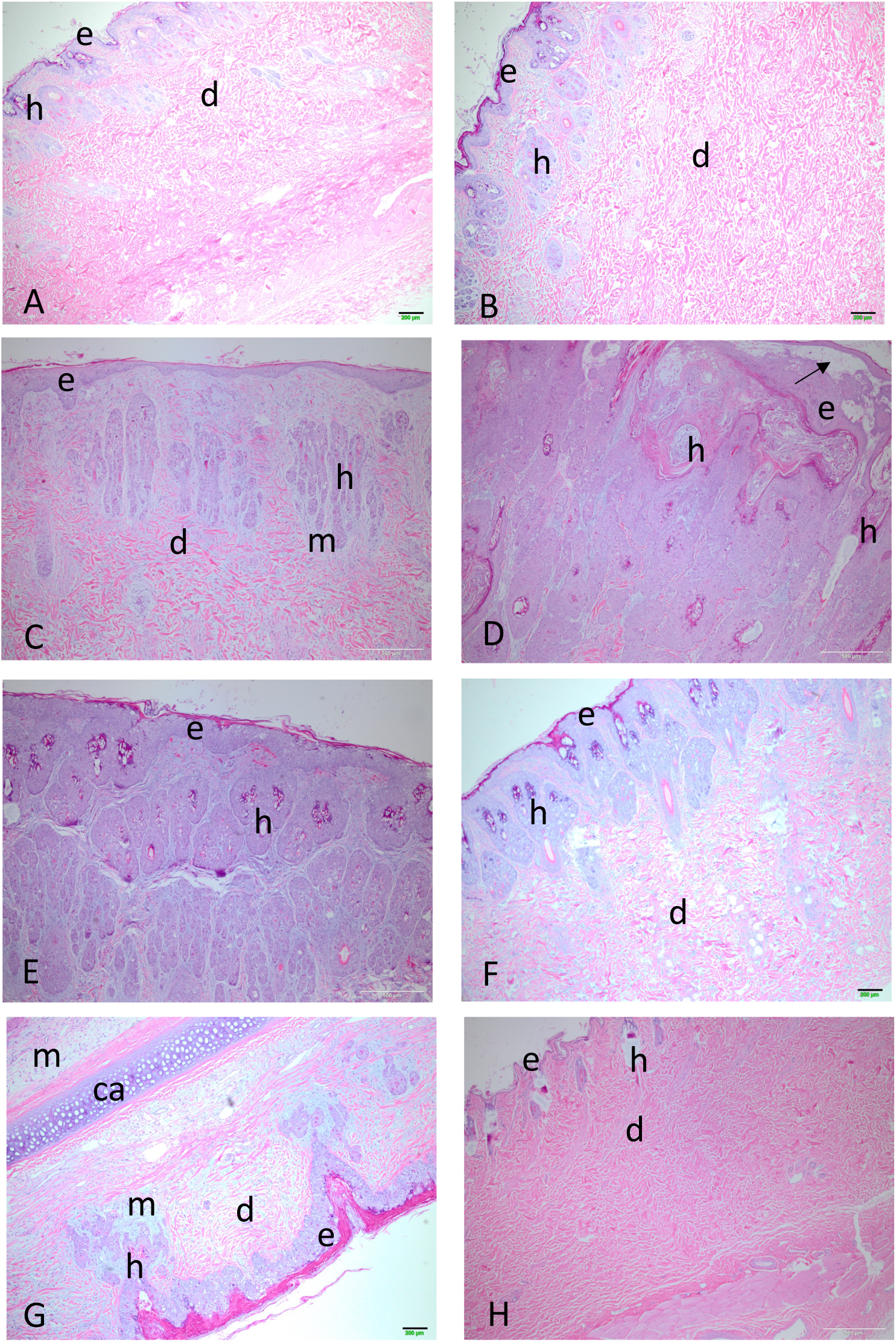
Primary lesion histology. (A) ABC Range (lineage B) day 12; e epidermis; d dermis; h hair follicle. (B) ABC Range day 19. (C) Mt Gambier (lineage C) day 11. (D) Mt Gambier day 21. (E) Wonga Park (lineage C) day 22. (F) Monarto (lineage A) day 18. (G) Section through ear Coomandook (lineage C) day 25; ca cartilage; k keratin; m mucin. (H) Full thickness section through normal skin from the inoculation site; e epidermis; d dermis; h hair follicle. All images are shown at the same magnification.

Fig. 7A and B show histology of the primary lesion from a rabbit infected with ABC Range (lineage b) at day 12 and another at day 19. There is very little disruption of the dermis at day 12 with the section showing a full thickness of the skin and even at day 19 the lesion is quite minimal although now there is some swelling although the dermal collagen fibres (pink stained) are still well organized. The lineage c viruses show much more proliferation of the epidermis, which replaces the upper dermis and considerable deposition of blue-grey-staining mucin-like material. For the lineage c virus Mt Gambier at day 11 (Fig. 7C) proliferation of epidermal cells seems to be from the hair follicles and there is already considerable deposition of mucin and loss of dermal architecture although pink-staining collagen bundles are still present lower down. At day 21 (Fig. 7D) the proliferation of epidermal cells has completely replaced the dermis and appears to be mostly arising from the hair follicles. There is vesiculation of the upper epidermis (arrow). For the lineage c Wonga Park at 22 days (Fig. 7E) this replacement of the upper dermis with epidermal cells is virtually complete but the outer layer of the epidermis is still intact (see also Supplemental material Fig. 1E). The lineage a virus Monarto at day 18 (Fig. 7F) is somewhat intermediate with proliferation of epidermal cells more pronounced than for the ABC Range virus but the structure of the lower dermis is reasonably well maintained. The histology in the secondary lesions and ears resembled that in the early-stage primary lesions with proliferation of epidermal cells, mucin deposition and minimal inflammatory cell response (Fig. 7G). For comparison, normal skin from the injection site is shown in Fig. 7H. Note the very thin epidermal layer and the organized collagen bundles of the dermis. Despite these histological differences between virus lineages, the actual virus titres within the lesions were generally similar (Fig. 5A) but the more extensive cutaneous proliferation of the base of the ears and head would have contained much more virus overall.

Pathology in the mucocutaneous tissues of the eyelids was similar to the primary lesions although sometimes more extreme, particularly in the epidermis, with very large vesicles often present; bacterial invasion and neutrophil infiltration could be seen in some late cases particularly below the mucosal epithelium and on the epidermal surface where, in contrast to primary lesions, thick scabbing occurred late (data not shown). Proliferation of epithelial cells from glandular elements was a feature of the eyelid pathology. As with the primary lesions there was a difference between time and virus lineage with earlier lesions showing much less

pathology and lineage c viruses tending towards more proliferative lesions, which could begin to resemble Lu but taking many days longer than the Lu lesions to develop. Even at 23 days the lineage b Acton 8-14 virus showed much less proliferative change than the lineage c viruses (data not shown).

## DISCUSSION

The aim of this study was to examine the phenotypic consequences of about 15 years of evolution of MYXV by characterising the disease caused by recent Australian MYXV isolates. A particular focus was whether there was any feature of the disease caused by the rapidly evolving lineage c viruses that might indicate ongoing adaptive evolution.

All of the Australian viruses tested induced acute death mimicking neutropenic septicaemia in some rabbits. Most had an amyxomatous phenotype with little or no inflammatory response in the cutaneous lesions. We hypothesise that the amyxomatous phenotype in laboratory rabbits is the result of selection against an inflammatory response in cutaneous lesions to enhance transmissibility in resistant wild rabbits. In some virus genetic backgrounds neutrophil activity is sufficiently suppressed that generalized bacteraemia can occur causing acute death in virtually all laboratory rabbits and mimicking very high grade I virulence but with a completely different pathogenesis to earlier grade I virulent viruses. In other genetic backgrounds some infected rabbits have a more prolonged disease course, and these viruses were of grade II to V virulence. The amyxomatous phenotype suggests that a neutrophil or other innate inflammatory response is involved in the formation of discrete, circumscribed primary and secondary lesions and the early scabbing that limits vector access in infections with viruses such as Lu and SLS. It is questionable whether the virulence exhibited by these modern viruses should be compared with the earlier viruses since the bacteraemia/toxaemia seems critical and we have previously demonstrated that survival time could be extended by treatment with antibacterials whereas antibacterials had no effect on SLS (8).

### Virulence and disease phenotypes in viruses from 2012-2015

Rabbits infected with viruses from each lineage developed the novel disease syndrome that we have previously described for Australian viruses from the 1990s and more recent viruses from Great Britain (8) (11). This was characterised by sudden collapse often with acute pulmonary oedema, and in many cases, bacteria distributed throughout multiple tissues and a lack of cellular inflammatory responses to the bacteria. The only predictive sign was that rectal temperatures were sometimes transiently elevated to ≥41°C in the 24 hours prior to death. As a working model, we interpret this as a direct or indirect suppression of neutrophil activity causing a syndrome that mimics neutropenic septicaemia (8). In contrast to the lack of neutrophils in these acute deaths, it has been shown elsewhere that laboratory rabbits infected with SLS had a peripheral blood neutrophilia late in infection. When tissues from laboratory rabbits infected with SLS are examined histologically, lymphocytes are depleted in virus-infected tissues such as lymph nodes or primary lesions, but neutrophils tend to be plentiful (8) (19). In the Lu infections described here, neutrophils were prominent in primary and secondary lesions, eyelids, lymph nodes, spleen, liver and lungs (Fig. 6g, h) at the same time points as acute deaths were occurring with the Australian viruses we tested. This supports the hypothesis that this acute disease phenotype is the result of one or more novel mutation(s) affecting neutrophil activity in non-resistant rabbits.

Importantly, when viruses were tested in the innately resistant CR line of laboratory rabbits, the same acute disease phenotype was observed in some rabbits in each group except those infected with the highly attenuated Wonga Park virus. This confirmed that the novel disease phenotype was independent of rabbit lineage. Acute pulmonary oedema had also been observed with viruses from the 1990s using outbred laboratory rabbits in Australia, but the mechanism and pathology of the disease syndrome had not been characterised (Kerr unpublished data). We have not yet been able to identify a specific viral mutation associated with this phenotype and it is possible that the mutation has a different mechanism of action in resistant wild rabbits.

All the Australian viruses tested, with the possible exception of TF2930 (which was only tested in CR rabbits), had the amyxomatous phenotype, which our data suggest is strongly linked to a lack of inflammation in the cutaneous lesions. All of the viruses were capable of inducing acute collapse/bacteraemia in at least some rabbits. However, some viruses, such as ABC Range were extremely lethal causing acute collapse in 6/6 Oak rabbits and 11/12 CR rabbits with short ASTs while others were much more attenuated with extended ASTs. At this stage we do not have a molecular explanation for this difference. However, ABC Range is the most virulent Australian virus we have tested and interestingly has reading frame disruptions in the *M005L/R* and *M153R* genes both of which have major virulence functions in the Lu virus (36) (37). It seems likely that the link between the acute death phenotype and the amyxomatous disease is the overall lack of an inflammatory response and that this implies that the inflammatory response, predominantly neutrophils, is important in development of discrete primary and secondary lesions early in infection. However, the amyxomatous disease and virulence seem poorly correlated suggesting that there are more factors involved that need to be identified.

### Does viral lineage c lead to differences in phenotype?

Lineage c MYXVs were isolated from a broad area of south-eastern Australia between 2012 and 2016 coincident with viruses from lineage a and b. Phylogenetically, we infer that a rapid, punctuated burst of evolution occurred in this lineage at some point between 1996 and 2012 (34). In contrast, there is no indication of a change in evolutionary rates in lineage a and b during this time.

We tested eight lineage c viruses in laboratory rabbits using viruses from different branches of the tree and from a range of geographic locations. Virulence grades ranged from I (Hattah, tested only in Oak rabbits) to IIIB/V (Wonga Park in Oak and CR rabbits respectively). Most were moderately attenuated grade II or III with a tendency to be more attenuated in the resistant CR rabbits. While most viruses tested had an amyxomatous phenotype, compared to the lineage a and b viruses tested, in the lineage c infections there appeared to be much more proliferative swelling at the base of the ears, around the eyelids and the skin of the anterior forelegs. In rabbits that survived for longer, this swelling became folded into firm ridges that could become distorted especially around the eyelids and base of the ears and eventually morph into more discrete lesions. This is a distinctly different pathogenesis from the secondary lesions that occur with the progenitor SLS, where widespread discrete secondary lesions develop from around six days after infection (19), and from the cutaneous lesions of Lu tested here. Virus titres in these swollen cutaneous tissues were extremely high and similar to titres in the primary lesions. In particular, the histology was very different to SLS and Lu with much more proliferation of epidermal cells that filled the dermis and an absence of the massive disruption of the dermis, and no evidence of small blood vessel disruption and no development of the large stellate “myxoma cells” that arise from the blood vessels in SLS and Lu cutaneous lesions and are often surrounded by neutrophils (23) (38).

Insect transmissibility is dependent on high titres of virus in cutaneous sites accessible to the vector. Studies on transmissibility have usually focussed on the primary lesion as a proxy for cutaneous titres. However, the virus-rich tissues of eyelids, face and ears are probably essential for transmission as mosquitoes are less likely to feed on the more densely haired parts of the body (20). This may be less important for flea transmission as rabbit fleas will move all over the body but still need to feed on areas of concentrated virus to contaminate their mouthparts for transmission. Scabbing of the cutaneous lesions as occurs with SLS and Lu limits vector access.

From the results obtained here, we hypothesise that lineage c viruses are capable of enhanced cutaneous dissemination to sites around the head where mosquitoes are more likely to feed and that they are able to suppress inflammatory responses at these sites allowing persistent virus replication to high titres. This does not seem to be necessarily linked to virulence at least in laboratory rabbits. In contrast, the lineage b viruses were generally of higher virulence with more limited cutaneous swelling around the head and ears although still with an amyxomatous phenotype. Although we have not tested enough recent lineage a viruses to draw any conclusions about these viruses, one CR rabbit infected with each lineage a virus (Monarto and BRK) developed the folded ears and face seen in the lineage c viruses suggesting that prolonged survival in resistant rabbits could enhance transmissibility. This indicates selection at multiple sites affecting both virulence and cutaneous immune suppression in the lineage c viruses. However, it is difficult to point to a genetic signature of this across the genetically diverged lineage c viruses. It seems likely that there has been selection against inflammatory responses and that in some virus genetic backgrounds this causes overwhelming immunosuppression with bacteraemia and acute collapse.

A feature of all lineage c viruses, except for Hoppers Crossing, was a G insertion at the 5’ end of the *M005L/R* gene (L/R indicates that the gene is present within the terminal inverted repeats and so there are two copies transcribed in opposite directions), which reverses a single nucleotide deletion present in the progenitor SLS virus and all subsequent Australian virus isolates sequenced. This deletion disrupted the *M005* reading frame with predicted loss of activity of any protein translated from this gene (34) (39). The M005 protein, also called T5, is an E3 ubiquitin ligase and has been associated with multiple functions in the Lu strain of MYXV where it is essential for replication in lymphocytes (40). Deletion of this gene from Lu leads to almost complete loss of virulence (36). Despite this virulence function in Lu, the lack of this gene appears not to have affected the virulence and spread of SLS and the viruses descended from SLS. Indeed, replacement of the disrupted M005 gene of the attenuated Uriarra (isolated in 1953) strain of MYXV with the Lu M005 had no effect on virulence (41). Loss of the active site of the M005 protein was also seen in highly virulent virus isolated from Yorkshire in Britain indicating that other mutations can compensate for the loss of M005 in the Lu background (11). It is interesting that early studies on the M005 protein in Lu indicated that its deletion prevented development of lesions at distal cutaneous sites (36).

In addition, in lineage c viruses, an A66V substitution occurred in the M010 protein, a viral homologue of epidermal growth factor, also associated with virulence (42). This is the only mutation we have seen in this conserved protein in Australian viruses. Whether this has any role in the extreme epidermal cell proliferation seen in rabbits infected with these viruses is not known. The equivalent gene in the closely related leporipoxvirus, Rabbit fibroma virus, also has valine at this position.

### Impact of rabbit strain

In the CR line of laboratory rabbits, average survival times tended to be longer and so virus virulence tended to shift towards more attenuated grades compared to the Oak line of rabbits. In addition, the outcomes for repeated measures of virulence could be inconsistent, skewed by the results of one or two rabbits. This is similar to the results seen for moderately attenuated (grade II/IIIA) viruses in non-resistant rabbits both here and in earlier work (8). Based on earlier work on resistance (19) (38) it is probable that the resistant CR rabbits had an enhanced innate immune response that was able to constrain virus-induced pathology. As with Australian wild rabbits, resistance could be overcome by viruses of high virulence (32) as seen here with ABC Range, which was considerably more virulent than Lu in the CR rabbits. It seems likely that a “resistance” allele was inadvertently fixed in the CR founder population at some stage by a genetic bottleneck, possibly due to a disease outbreak or perhaps simply a stochastic event such as selection of a small pool of foundation breeding stock.

Fenner and Woodroofe (1965) (27) noted that when local (?) laboratory rabbit lines used for virulence testing in Australia were changed to NZ White rabbits a longer average survival time was observed with SLS (10.8 vs 12.5 days) and occasional NZ White rabbits infected with an otherwise lethal virus had an accelerated recovery. This occasional recovery suggests that resistance alleles were present in the outbred NZ White laboratory rabbit population but rare. Attempts to select laboratory rabbits for resistance to myxomatosis met with limited success (43).

We also saw occasional accelerated recoveries in the resistant rabbit trials. Of particular interest, these rabbits tended to develop widespread discrete secondary cutaneous lesions early in the disease course although the overall disease could be quite mild and was classified as category D, although the disease was quite different to that caused by attenuated viruses from the 1950s ((8). The remaining rabbits within the group developed an amyxomatous (category B or C) disease. As already discussed, this suggests that the formation of discrete secondary lesions is more likely to occur when there is some degree of innate and/or adaptive immune response to the virus infection.

### Ongoing evolution of virulence and resistance

The main measure of MYXV evolution in Australia from 1950 through to 1981 was testing virus isolates for virulence using the Fenner grading system in laboratory rabbits. Through most of this period the virulence of field isolates was relatively stable with most isolates being of intermediate grade III or grade IV virulence and small numbers with higher and lower virulence (17). There may have been a trend to increased numbers of grade II viruses in some areas associated with high resistance (44) (17). The trend to higher virulence is supported by the apparent increase in resistance found from wild rabbits in the late 1970s, particularly enhanced survival to challenge with the Lu strain, which in resistant rabbits is more virulent than SLS (45). When tested in the 1990s, wild rabbits infected with SLS mostly made an uneventful recovery indicating very high resistance (19) (31).

Something appears to have changed in this evolutionary dynamic between 1980 and the early 1990s. Detailed genetic and phenotypic studies of viruses from the 1990s in laboratory rabbits showed that the disease caused in laboratory rabbits was quite different to earlier myxomatosis (8). When tested in contemporaneous wild rabbits, the same viruses had a prolonged incubation period with little or no primary lesion visible at the inoculation site and caused more typical swelling around head, ears and eyelids, from which virus could be transmitted for prolonged times, and case fatality rates of around 50% or less (46). This suggests that in the field these viruses were effectively grade IV or V virulence. The change in disease phenotype seen in laboratory rabbits strongly supports a model where mutations in previously attenuated viruses have caused alterations in disease phenotype rather than a re-emergence of virulent viruses that had continued to circulate undetected for some 40 years (39). This timeline and selection by increased resistance is further supported by similar disease phenotypes occurring in viruses from Australia and Great Britain. Strong resistance emerged in the 1970s in Britain (28) (29) potentially driving virus adaptation. However, it has not proved possible so far to identify specific mutations associated with this return to virulence in viruses from either country. Since the novel disease phenotype was found in viruses isolated in Australia and Britain, it seems probable that this was an evolutionary convergent response to increasing resistance in the rabbit population – a biological “arms race”.

The continued isolation of highly virulent, attenuated and moderately attenuated viruses (as measured in laboratory rabbits) suggests that under different ecological conditions or in different rabbit populations viruses with quite different virulence are able to establish and transmit. Does this mean that the nexus between virulence and transmission is breaking down? Earlier studies on flea transmission demonstrated that rabbits that survived infection were always poor sources of infection as were those that died early (47). This suggested a strong correlation with moderate virulence and transmissibility. Studies with 1990s viruses in highly resistant wild rabbits showed that most did not induce a significant primary lesion and that systemic spread to secondary cutaneous sites such as eyelids and base of the ears was likely critical for transmission and that virus could persist at these sites for prolonged periods (46). In its natural hosts MYXV and the related rabbit fibroma virus cause cutaneous fibromas limited to the inoculation site (at least in immunocompetent rabbits). Whether MYXV and European rabbits can co-evolve to such a “climax” state probably depends on whether systemic spread and multiple cutaneous lesions leads to more overall transmission, better fitness, than a single fibroma like lesion and whether European rabbit populations have immune system alleles that could constrain virus replication to the initial cutaneous site. If dissemination is critical for transmission, then mutations in the virus that favour dissemination by suppressing immune responses may also increase virulence.

In Europe, a recent recombination event between MYXV and an unknown poxvirus has led to a virus that can transmit and cause disease in both rabbits and hares (48), which may lead to new evolutionary pathways.

## METHODS

### Rabbits

All animal experiments were approved by The Pennsylvania State University Institutional Animal Care and Usage Committee: permit numbers: 33615 and 42748.

Male New Zealand White rabbits 4 months of age were obtained from either Harland Laboratories, Oakwood facility, Pa, USA (“Oak rabbits”) or Charles River, Canada (“CR rabbits”) and were free of multiple pathogens including *Pasteurella multocida* and *Bordetella bronchiseptica*. Rabbits were housed as previously described (11), in individual cages in temperature-controlled rooms on a 12 hour light/dark cycle and fed on standard rabbit pellets. All animals were allowed at least 10 days to acclimate prior to experimental infection. Food consumption was recorded each day and water intake estimated. Inappetent animals were supplemented with hay and vegetable-based baby food. Faecal and urinary outputs were monitored by inspection of the paper sheets lining the litter trays.

### Virus phenotype assays

Rabbit trials were conducted as previously described (11). For each of 16 viruses tested, six rabbits were inoculated intradermally over the rump with 100 pfu of virus. Trial 1 tested five viruses in Oak rabbits; Trial 2 tested 10 viruses in CR rabbits and Trial 3 compared 7 of these viruses head-to-head in CR and Oak rabbits plus the Lausanne strain (Lu) as a virulent control. In each trial, animals were clinically examined in the morning and rectal temperature and body weight measured. A semi-quantitative score of 0 (normal) to 3 (severe) was given for each of demeanour (encompassing activity, muscle strength, alertness), eyelid swelling, ano-genital swelling, ear swelling, respiratory difficulty and scrotal involvement. Primary lesions were measured and described, and distribution and size of secondary lesions recorded as well as other clinical signs such as head swelling, bruising/haemorrhages, diarrhoea, nasal and ocular discharges, muscle fasciculations, ataxia etc. Once systemic signs of disease became apparent, rabbits were also monitored in the afternoon. Animals that died or were euthanised were necropsied as soon as possible after death and refrigerated until necropsy. Criteria for euthanasia were as previously described (11). Note that in trial 3 only five rabbits were infected in the Oak group for Wonga Park virus because one rabbit assigned to this group acclimated poorly to feeding and so was not used for infection. The disease caused by the Lu virus is much more florid in terms of head, eyelid, ear and ano-genital swelling that the 0-3 scoring used for the Australian viruses was not comparable for Lu.

### Survival times

Survival times for animals that died unobserved were assigned as the midpoint between observations unless there was clear evidence that death had occurred in the previous one to two hours (warm body and lack of rigor mortis). For the purpose of estimating AST and Kaplan-Meier survival analysis, animals that were euthanized when moribund or becoming moribund were assigned a survival time based on estimated likely time to death. These survival times were likely accurate to within hours. Animals euthanized on humanitarian grounds were assigned an additional survival time of 24-72 hours based on estimated likely time to death. These survival times may be underestimates. For estimating AST, rabbits classified as likely survivors were assigned a survival time of 60 days (5). For assignment to virulence grades, AST was calculated as the mean of survival times that were transformed as log_10_(ST-8) as per (5) and then back-transformed. Criteria for virulence grades are shown in Table 1. Kaplan-Meier survival analyses were done in Sigmaplot and used the individual estimated survival times not the transformed times.

For virus titres in tissues and for disease category assignments (see below) the actual time of death or euthanasia was used without adjustment, other than for unobserved deaths where midpoint time was used as above and is referred to as actual survival time.

### Disease appearance classification

In earlier work, where we tested viruses from the 1950s and the 1990s, we assigned clinical appearance to four categories: (A) Typical cutaneous nodular myxomatosis; (B) Acute collapse with few overt signs of myxomatosis; (C) Progressive amyxomatous disease in rabbits infected with viruses causing category B but that did not suffer acute collapse within 15 days; and (D) Attenuated cutaneous nodular myxomatosis (8). Here we have classified clinical appearance using this scheme but only the Lu virus caused typical cutaneous nodular myxomatosis (category A) and none of the viruses consistently caused attenuated cutaneous myxomatosis (category D). However, some acute deaths occurred with much more swelling so that category B could have been divided into B1 and B2 and similarly some viruses tended to cause a much more progressive amyxomatous disease than others with much more swelling and folding of the skin of the head and swelling of the eyelids and hence category C could have been further subdivided into C1 and C2.

Typical cases for each category are shown in Fig. 1A.

### Sample collection

Sample collection and processing for histology were as previously described (11). For virus titrations, homogenized tissue samples were plaque-assayed on RK-13 cell monolayers. Samples for histology and virus titration were not collected from rabbits that died overnight due to the advance autolysis that occurred.

### Viruses

The viruses tested are listed in Table 2. Apart from the Lausanne strain (Lu), all viruses were isolated in Australia. The Lu stock was prepared from virus originally produced by the Commonwealth Serum Laboratory for field release in Australia. Viruses were isolated in RK-13 cells and working stocks amplified, and titred by plaque assay, in RK-13 cells as previously described (39). The complete genome sequence was determined for each virus (34, 39).

**TABLE 2.** Viruses tested. Shaded viruses were used as controls.

| **Virus (lineage)** | **Virus name used in text** |
| --- | --- |
| Aust/Vic/Wonga Park/03-2012 (c)<sup>1</sup> | Wonga Park |
| Aust/SA/Turretfield(1213)/10-2012 (c) | TF1213 |
| Aust/Vic/Hattah/10-2012 (c) | Hattah |
| Aust/SA/ Turretfield(2930)/07-2012 (c) | TF2930 |
| Aust/SA/Adelaide Hills/05-2012 (c) | Adelaide Hills |
| Aust/SA/Mt Gambier/01-2015 (c) | Mt Gambier |
| Aust/SA/Coomandook/12-2013 (c) | Coomandook |
| Aust/Vic/Hoppers Crossing/03-2012 (c) | Hoppers Crossing |
| Aust/NSW/Euchareena(3)/2-2013 (b) | Euchareena |
| Aust/SA/ABC Range/12-2014 (b) | ABC Range |
| Aust/ACT/Acton/12-2012 (b) | Acton-12 |
| Aust/ACT/Mulligans Flat/04-2013/1 (b) | Mulligans Flat |
| Aust/ACT/Acton/08-2014 (b) | Acton 8-14 |
| Aust/SA/Monarto Zoo/11-2013 (a) | Monarto |
| Aust/NSW/Brooklands/4-93 (a) | BRK |
| Brazil/Campinas/1949 Lausanne strain | Lu |

<sup>1</sup>Nomenclature: Australia/State/location (sample identifier)/month-year/isolate number.

**TABLE 3.** Summary of trial 2 and 3 virulence data.

| **Virus** | **Trial 3**<br>**AST<sup>1</sup> Oak** | **Trial 3**<br>**AST CR** | **Trial 3**<br>**CFR<sup>2</sup> Oak** | **Trial 3**<br>**CFR CR** | **Virulence**<br>**grade Oak** | **Virulence**<br>**grade CR** | **Virulence**<br>**grade trial 2** | **AST trial 2** |
| --- | --- | --- | --- | --- | --- | --- | --- | --- |
| Lausanne | 12.5 | 13.2 | 6\6 | 6\6 | I | II | <sup>3</sup>nd | nd |
| ABC Range (b) | 9.1 | 11.4 | 6\6 | 6\6 | I | I | I | 12 |
| Acton 8-14 (b) | 12.7 | 11.8 | 6\6 | 6\6 | I | I | IIIA | 19.1 |
| Adelaide Hills (c) | 14 | 13.3 | 6\6 | 6\6 | II | II | II | 13.2 |
| Hoppers Crossing<br>(c) | 13.1 | 17 | 6\6 | 6\6 | II | IIIA | II | 13.8 |
| Mt Gambier (c) | 14.6 | 16.5 | 6\6 | 6\6 | II | IIIA | IIIA | 21.4 |
| Coomandook (c) | 16.3 | 24 | 6\6 | 5\6 | IIIA | IIIB | IIIB | 25.9 |
| Wonga Park (c) | 23 | n/a | 5\6 | 0\6 | IIIB | V | nd | nd |

<sup>1</sup>AST: average survival time was calculated using the transformation of Fenner and Marshall (1957) where survival time is transformed to log<sub>10</sub>(ST-8) and the transformed values used to calculate average survival times and then back-transformed.
<sup>2</sup>CFR – case fatality rate
<sup>3</sup>nd – not done.

