## Supplemental Figures for "Divergent Evolutionary Pathways of Myxoma Virus in Australia: Virulence Phenotypes in Susceptible and Partially Resistant Rabbits Indicate Possible Selection for Transmissibility"

**Fig S1**. (A) Wild rabbit from which Acton-12 was isolated: coalescing cutaneous lesions under the chin (arrow). (B) Swollen eyelids (white arrow); discrete secondary lesions in ear (black arrow). (C) Wonga Park (Oak rabbit) day 16 after infection. Extreme swelling of ears and eyelids but eyelids are still open. (D) Wonga Park (Oak rabbit day 22 autopsy). Looking down on head (arrow indicates swollen, closed left eyelid). Extreme swelling and folding of skin over face and folding of ears. (E) Primary lesion of same rabbit (arrow) showing minimal response at inoculation site. (F) Primary lesion (long arrow) from a Lu infected Oak rabbit (day 10); short arrow indicates beginning of scabbing at edge of lesion. Note the raised, crimson, demarcated lesion compared to the Wonga Park primary lesion.

**Fig S2**. Scatter plots of titres at actual time of death for each rabbit. A. Popliteal lymph node. B. Spleen. (C) Lung. D. Liver. Regression lines with 95% confidence intervals are plotted.

**Fig S3**. Boxplots of titres at actual time of death for each virus. (A) Popliteal LN. (B) Spleen. (C) Lung. (D) Liver. (E) Primary lesion.

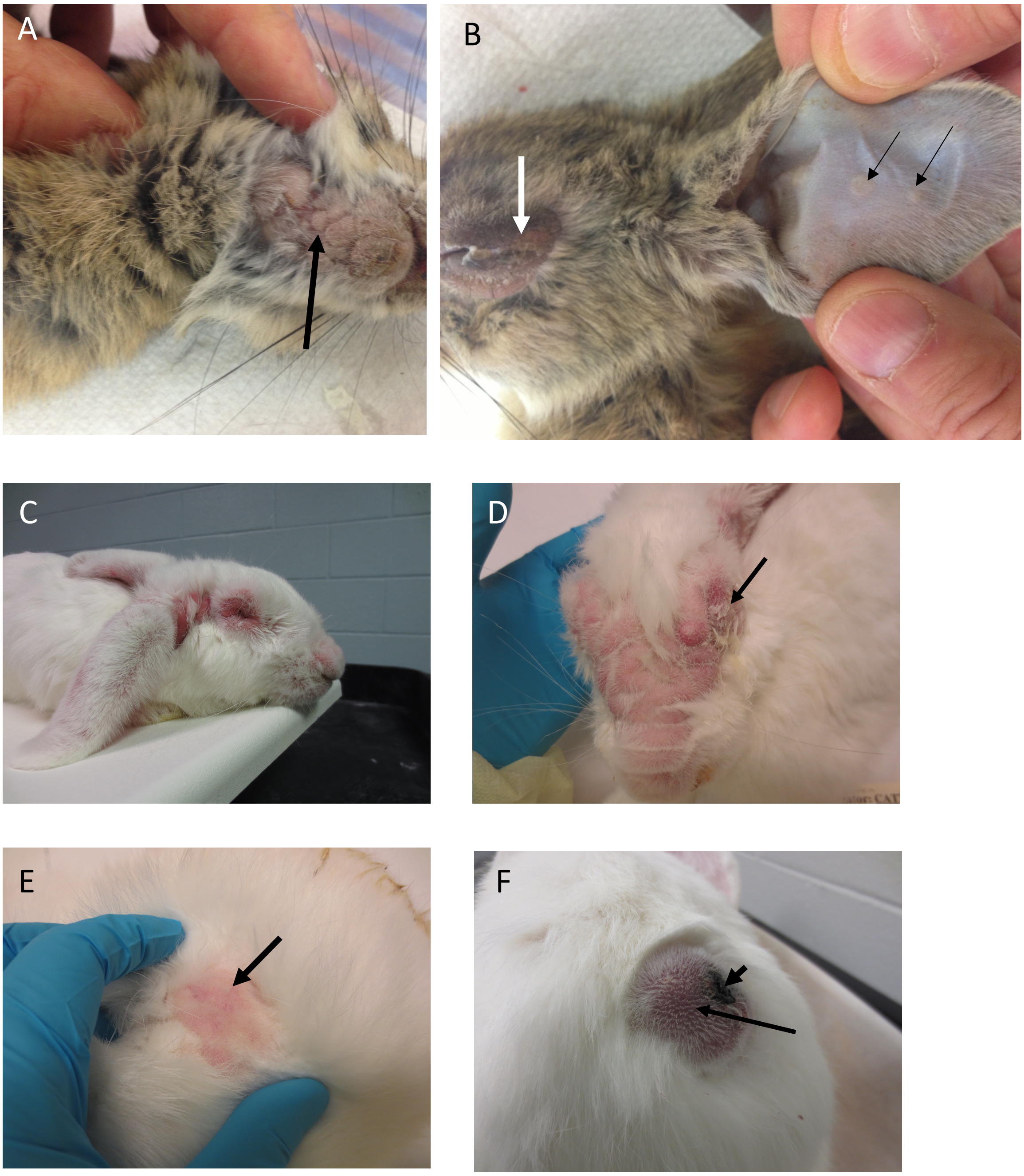


**FIG S1.**

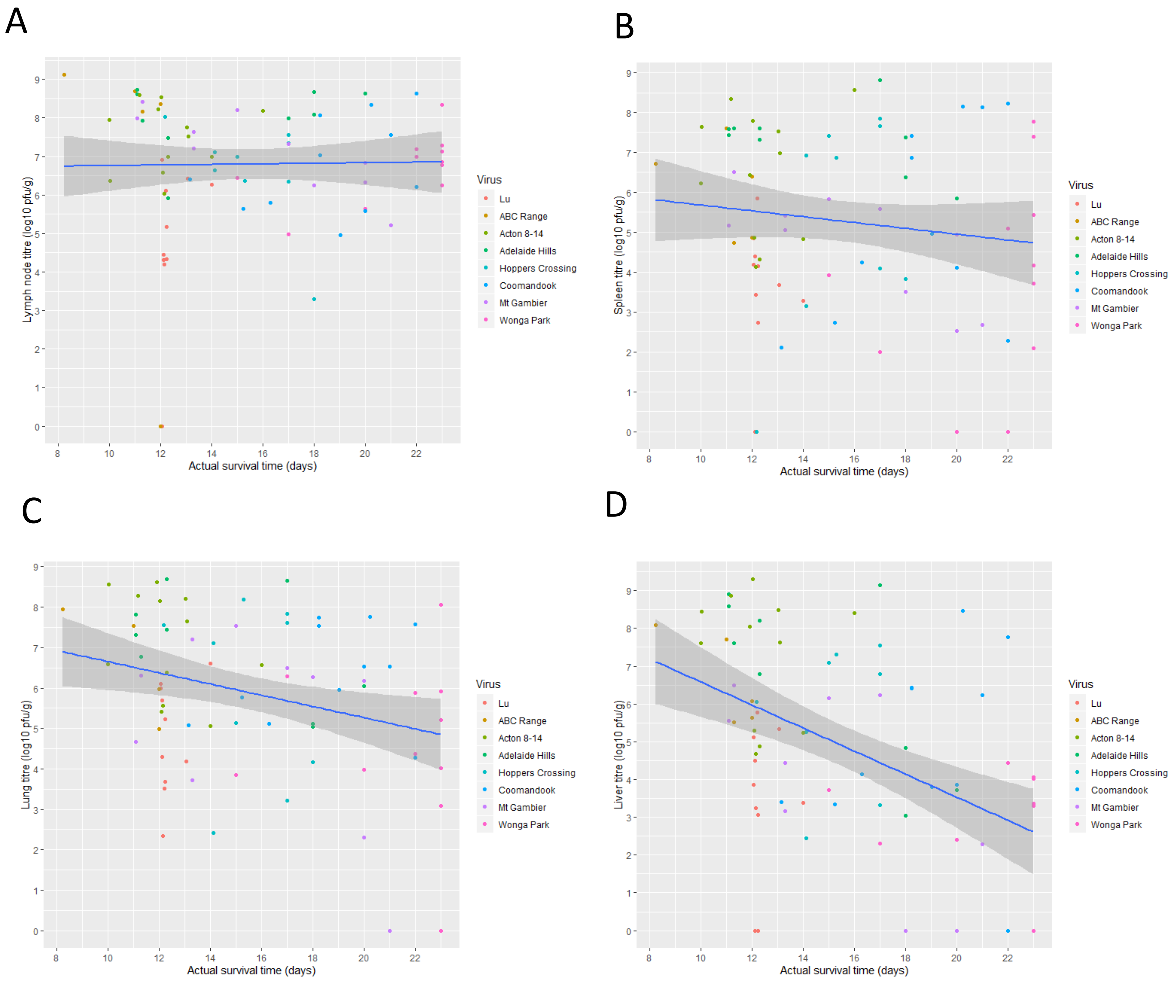
A
B
C
D
Lymph node titre (log10 pfu/g)
Spleen titre (log10 pfu/g)
Lung titre (log10 pfu/g)
Liver titre (log10 pfu/g)
Actual survival time (days)
Virus
Lu
ABC Range
Acton 8-14
Adelaide Hills
Hoppers Crossing
Coomandook
Mt Gambier
Wonga Park

**FIG S2.**

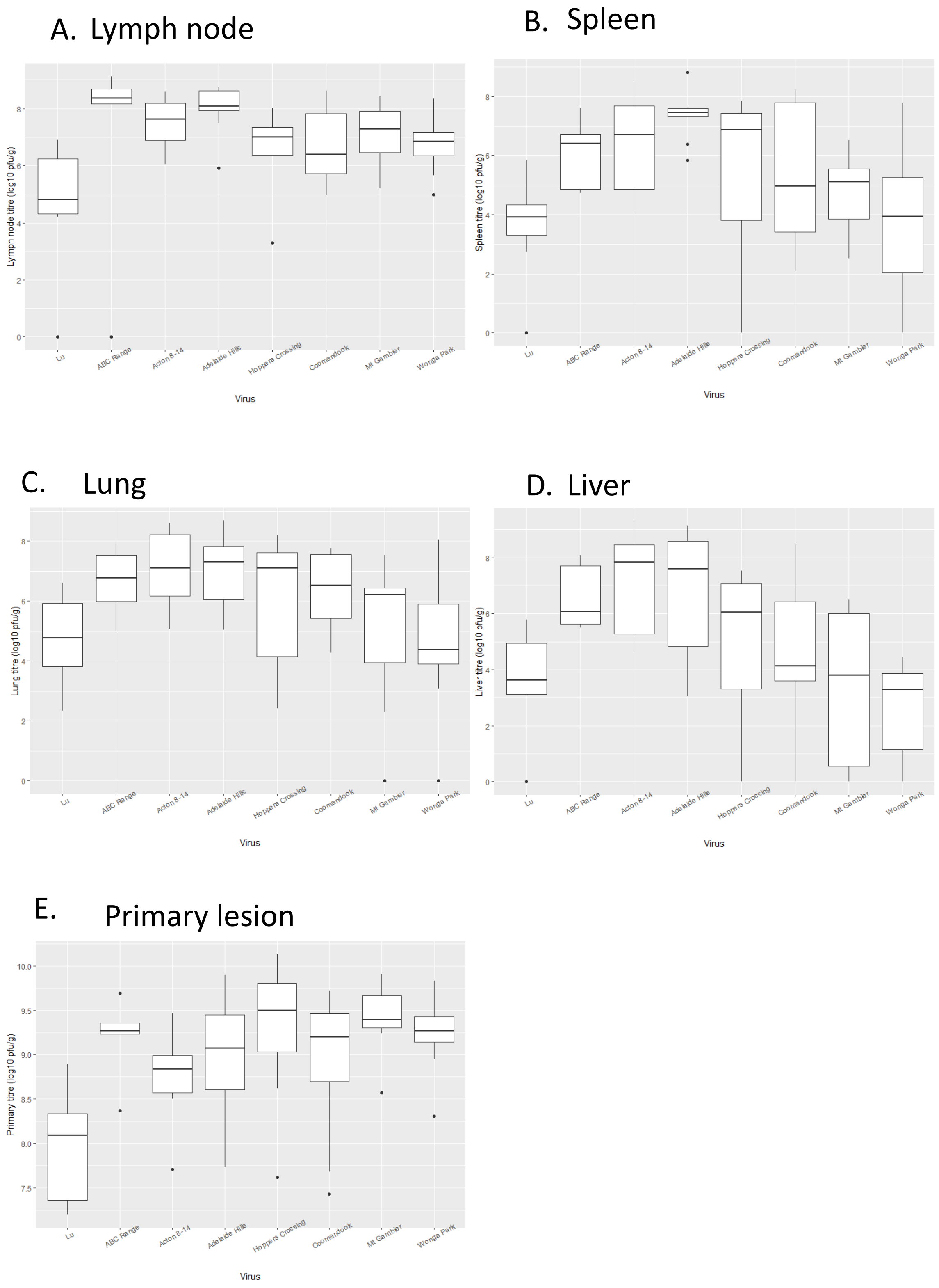
A. Lymph node
Lymph node titre (log10 pfu/g)
B. Spleen
Spleen titre (log10 pfu/g)
C. Lung
Lung titre (log10 pfu/g)
D. Liver
Liver titre (log10 pfu/g)
E. Primary lesion
Primary titre (log10 pfu/g)
Lu
ABC Range
Acton 8-14
Adelaide Hills
Hoppers Crossing
Coomandook
Mt Gambier
Wonga Park
Virus

**FIG S3.**
